# Pathogenic potential of Hic1 expressing cardiac stromal progenitors

**DOI:** 10.1101/544403

**Authors:** H Soliman, B Paylor, W Scott, DR Lemos, CK Chang, M Arostegui, M Low, C Lee, D Fiore, P Braghetta, V Pospichalova, CE Barkauskas, V Korinek, A Rampazzo, K MacLeod, TM Underhill, FM Rossi

## Abstract

The cardiac stroma contains multipotent mesenchymal progenitors. However, lineage relationships within cardiac stromal cells are still poorly understood. Here, we identify heart-resident PDGFRa^+^ Sca-1^+^ cells as cardiac Fibro/Adipogenic Progenitors (cFAPs) and show that they respond to ischemic damage by generating fibrogenic cells. Pharmacological blockade of this differentiation step with an anti-fibrotic tyrosine kinase inhibitor decreases post-myocardial infarction (MI) remodeling and leads to improvements in heart function. In the undamaged heart, activation of cFAPs through lineage-specific deletion of the quiescence factor Hic1 reveals additional pathogenic potential, causing fibro-fatty infiltration of the myocardium and driving major pathological features of Arrhythmogenic Cardiomyopathy (AC).

**Highlights:**

- A subpopulation of PDGFRa^+^, Sca-1^+^ cells, previously considered to be a sub-type of cardiac fibroblasts, are multipotent mesenchymal progenitors,
- Cardiac damage triggers the differentiation of PDGFRa^+^ Sca-1^+^ cells into Sca-1^-^ cells expressing a fibrogenic transcriptional programme,
- Blockade of the cFAP-to-fibroblast transition by Nilotinib ameliorated cardiac dysfunction post-MI and modulated cardiac remodelling.
- Studies performed on a model of experimentally-induced AC confirmed that cFAPs are a source of both cardiac fibroblasts and adipocytes *in vivo*.
- Conversely, in the undamaged heart, activation of cFAPs by means of lineage-specific deletion of transcription factor Hic1, resulted in fibro/fatty cardiac degeneration and pathological alterations reminiscent of AC. Collectively, our findings show that a proportion of what are commonly termed “fibroblasts” are actually multipotent mesenchymal progenitors that contribute to different forms of cardiac degeneration depending on the damage setting.

---

The cardiac stroma plays important roles both in homeostasis and remodelling following damage. In the aftermath of myocardial infarction, stromal cells activate a fibrogenic programme that eventually leads to the replacement of lost cardiac muscle with a rigid fibrous scar(Frangogiannis, 2012). In this case, fibrogenic cells take on a reparative role, as scar formation is critical to maintain appropriate cardiac function. Indeed, inhibition of this process by blocking TGFβ1 signalling results in increased post-MI mortality(Frantz et al., 2008; IKEUCHI et al., 2004). However, chronic interstitial fibrosis is also a common outcome of post-MI myocardial remodelling and is associated with multiple disease settings, suggesting that persistent activation of a fibrogenic program can also contribute to cardiac pathology(See et al., 2005). Despite the importance of cardiac fibrogenesis, the cellular and molecular mechanisms leading to accumulation of fibrogenic cells within the heart are poorly understood, limiting our ability to modulate them therapeutically.

Tissue resident cardiac stromal cells have been shown to give rise to multiple lineages including fibrogenic cells, committed osteoprogenitors and adipocytes(Chong et al., 2011). Those findings suggested that a mesenchymal hierarchy exists within the adult heart, comprising immature progenitors that give rise to more differentiated cell types. In particular, the circumstances under which progenitors generate the cells responsible for the pathological deposition of extracellular matrix remain poorly understood. Cardiac fibroblasts, defined based on the expression of classical markers such as collagen 1a1, TCF21 and PDGFRa, are heterogeneous and encompass several phenotypically and functionally distinct cellular subsets (Pinto et al., 2016). This lack of uniformity suggests that mesenchymal progenitors may have been included in such definition. Indeed, significant phenotypic similarities between classical cardiac fibroblasts and mesenchymal progenitors have been reported (Haniffa et al., 2009; Hematti, 2012).

Others and us have identified a population of tissue-resident multipotent stromal cells in skeletal muscle that is characterized by the expression of PDGFRa and the stem cell marker Sca-1(Joe et al., 2010; Uezumi et al., 2010). These cells are referred to as Fibro Adipogenic Progenitors (FAPs) due to their spontaneous ability to generate fibroblasts as well as adipocytes in vivo and in vitro. In addition, they can be induced to acquire an osteogenic fate upon treatment with BMP2, further supporting the notion that FAPs are multipotent progenitors. Phenotypically identical cells with similar developmental potential were also found in numerous other tissues, including bone, kidney, skin(Festa et al., 2011), fat(Chun et al., 2013) and the heart(Chong et al., 2011). These populations have been demonstrated to possess highly similar transcriptomes, albeit with some tissue-specific differences in gene expression patterns(Pelekanos et al., 2012).

Interestingly, unlike in skeletal muscle where all PDGFRa^+^ cells express the stem cell marker Sca-1, in the adult heart at steady state, a subpopulation of PDGFRa^+^ cells are Sca-1^-^. These Sca-1^-^ cells are likely to represent a terminally differentiated population as they have limited proliferative capacity and they do not clonally expand in culture(Chong et al., 2011).

Here we show that a surprisingly large fraction of stromal cells previously thought to be cardiac fibroblasts are in fact multipotent progenitors, as defined by their in vivo and in vitro lineage capabilities. We find that these progenitors generate PDGFRa^+^, Sca-1^-^ fibroblasts upon injury, and pharmacological inhibition of this differentiation step improves cardiac function following myocardial infarction. Of note, we also report that impairing the ability of these cells to remain quiescent by deletion of transcription factor Hic1 leads to fibrofatty infiltration and arrhythmias in the absence of primary cardiomyocyte damage. Consistent with this, lowering the threshold of activation of cFAPs in a model of arrhythmogenic cardiomyopathy (AC) leads to earlier onset and increased severity of the pathology.

## Results

### Hic1 expression identifies a population of cardiac stromal cells enriched in multipotent mesenchymal progenitors

Hic1 has been recently proposed as a transcription factor expressed in stromal progenitor cells (Underhill, ISSCR meeting 2014). In order to analyze the population of Hic1-expressing cardiac cells within the heart and gain a better understanding of their lineage capabilities *in vivo*, we took advantage of a double transgenic mouse carrying both tamoxifen (TAM) inducible Cre recombinase inserted in the Hic1 locus and the floxed tdTomato reporter (Hic1Cre^ERT2^/tdTomato; Fig. 1A). TAM treatment in these mice leads to indelible labeling of Hic1-expressing cells and their progeny with tdTomato, allowing lineage tracing. One week after TAM treatment, hearts were dissected and enzymatically dissociated. Td-Tomato^+^ cells were sorted and analyzed by single cell RNA sequencing (scRNAseq) to assess their degree of lineage commitment and establish population heterogeneity. Principal component analysis followed by dimensionality reduction revealed two spatially distinct clusters of cells, of which the largest represented PDGFRa^+^ stromal cells (Fig. 1B (left) and Suppl. Fig. S1A), and the other an as yet uncharacterized PDGFRa^-^, Sca-1^-^ cell type expressing pericytic markers that will be the subject of future study (Suppl. Fig. S1B).

**Figure 1.**
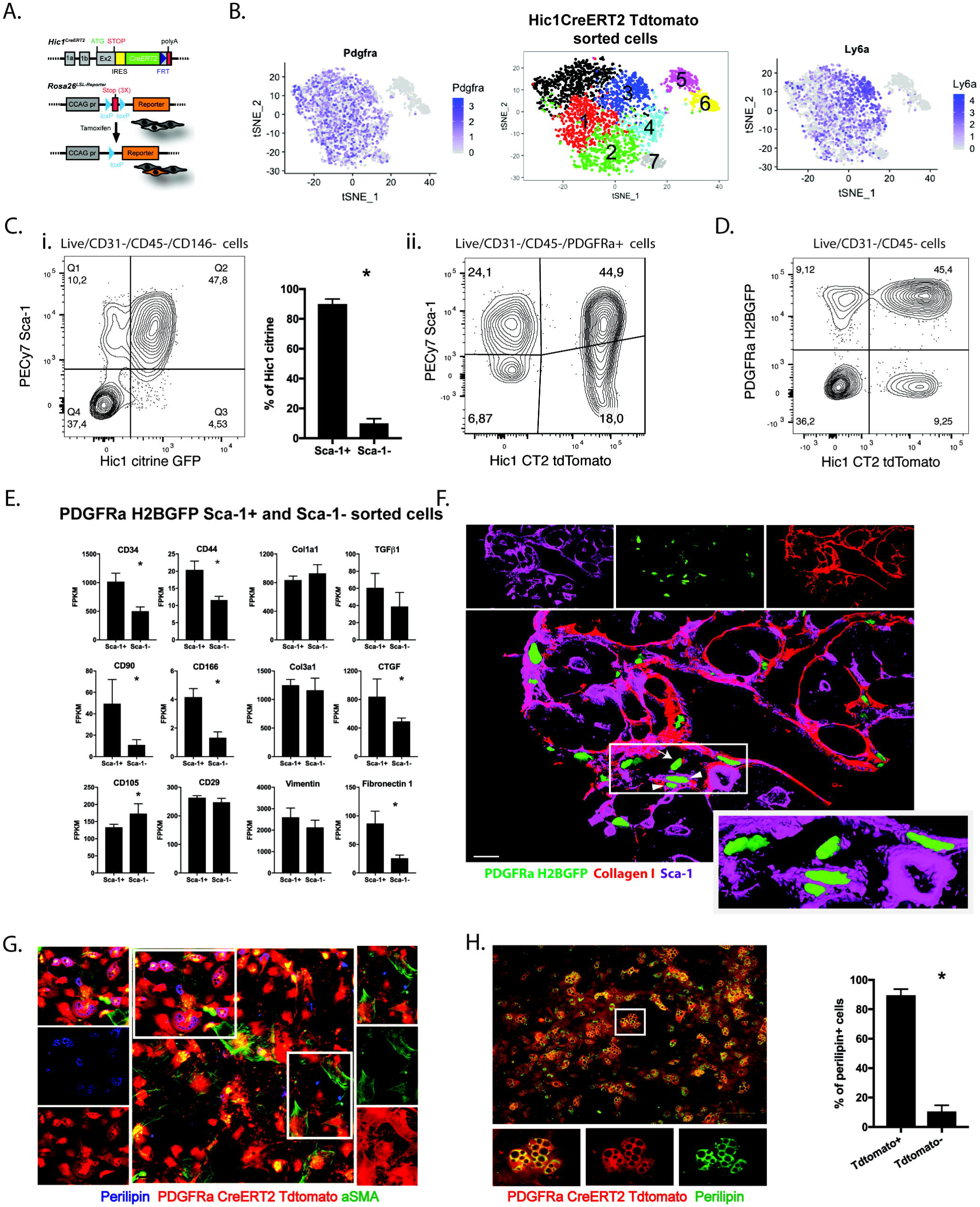
The PDGFRa^+^ Sca-1^+^ subset of cardiac fibroblast contains multipotent mesenchymal progenitors. A. Schematic of the Hic1Cre^ERT2^ and Rosa26^LSL-Reporter^ alleles. TAM administration leads to the Hic1-directed expression of the reporter tdTomato. B. t-SNE clustering plots of single cell RNA sequenced Hic1Cre^ERT2^/tdTomato^+^ cells in the heart showing pdgfra (top) and ly6a (Sca-1; bottom) RNA expression. C. Representative FACS plot and quantification of the CD45^-^ CD31^-^ CD146^-^ Sca-1^+^ and Sca-1^-^ cells in the heart that are Hic1 citrine GFP^+^ (i). (ii) Representative FACS plot of CD45^-^ CD31^-^ PDGFRa^+^ Sca-1^+^ and Sca-1^-^ that were Hic1 Cre^ERT2^/tdTomato^+^ (n=3 in each group). D. Representative FACS plot of CD45^-^ CD31^-^ PDGFRa H2BGFP^+^ and Hic1Cre^ERT2^/tdTomato^+^ cells in the heart. E. Quantification of RNA expression levels of the mentioned genes in CD45^-^ CD31^-^ PDGFRa H2BGFP^+^ Sca-1^+^ and Sca-1^-^ cells in the undamaged heart (n=4 for the Sca-1^-^ group and 5 for the Sca-1^+^ group). F. Representative 3D reconstruction of a z-stack of images of collagen I (red) and Sca-1 (magenta) immunostaining of 10µm cryosections from PDGFRa H2BGFP mice hearts. Inset: increased magnification of the area highlighted by the white rectangle showing PDGFRa H2BGFP and Sca-1 staining only. Arrowheads indicate examples of Sca-1^+^ cells and arrows indicate examples of Sca-1^-^ cells. Scale bar = 20µm. G. Representative images of perilipin (blue) and a-smooth muscle actin (a-SMA; green) immunostaining of fixed and permeabilized cells isolated from PDGFRa Cre^ERT2^/tdTomato mice hearts and single cell cultured in 96 well plates Scale bar=100 pixels. H. Representative image and quantification of perilipin^+^ PDGFRa Cre^ERT2^/tdTomato^+^ and PDGFRa Cre^ERT2^/tdTomato^-^ adipocytes (n=4 in each group). Scale bar=100µm. Data represented as mean ± SEM, *p<0.05.

Expression of PDGFRa is commonly used to identify stromal cells in the heart and other tissues(Joe et al., 2010; Uezumi et al., 2010). However, this population is highly heterogeneous, and likely to comprise multiple functionally different subsets. Indeed, cluster analysis of the PDGFRa+ cells revealed 6 different subclusters (Fig. 1B middle). Interestingly, cells expressing highest levels of the progenitor cell marker Sca-1 (Fig. 1B right) were assigned to the same subcluster (subcluster 3 shown in Fig. 1B and Suppl. Fig. S1C), suggesting that expression of this marker may identify a cell type expressing a distinctive transcriptional programme within the Hic1 Cre^ERT2^ tdTomato^+^ PDGFRa^+^ cells. Suppl. Fig. S1D shows a heatmap of the top 10 most differentially expressed genes in each of the PDGFRa^+^ subclusters.

To further investigate the relationship between Sca-1 and Hic1 expression we compared cells purified from hearts of Hic1Cre^ERT2^/tdTomato mice and from mice carrying citrine GFP knocked into the Hic1 locus (Hic1-citrine), which labels only cells expressing Hic1 and not their progeny (Pospichalova et al., 2011). We found that while Hic1 expression, after excluding pericytes, was mostly restricted to cells expressing Sca-1 (Fig. 1C, i), their tdTomato^+^ progeny included a fraction of Sca-1^-^ cells (Fig. 1C, ii). Based on our previous studies in the skeletal muscle(Joe et al., 2010; Uezumi et al., 2010), we hypothesized that Hic1^+^ PDGFRa^+^ Sca-1^+^ cells may be endowed with progenitor properties and that more differentiated Hic1^-^ PDGFRa^+^ Sca-1^-^ cells may represent their progeny.

To further assess differences between PDGFRa^+^ Sca-1^+^ and Sca-1^-^ cells, we sorted each subset from the stromal fraction (CD45^-^/CD31^-^/PDGFRa^+^) of enzymatically dissociated hearts. Initial analysis using digital droplet PCR showed that, compared to CD45^-^/CD31^-^ /PDGFRa^-^ cells, both PDGFRa^+^ populations were enriched for ECM^-^associated genes including TGFβ1, col1a1, vimentin, fibronectin-1 and connective tissue growth factor (CTGF) confirming their stromal nature(Ieronimakis et al., 2013) (Suppl. Fig. S1E). Deeper transcriptional analysis of the Sca-1^+^ and Sca-1^-^ populations using bulk RNAseq showed that the Sca-1^+^ population expressed significantly higher levels of progenitor markers including CD90, CD44 and CD34 (Fig. 1E). In contrast to previous reports(Ieronimakis et al., 2013) however, we failed to detect distinct anatomical locations for cardiac PDGFRa^+^/Sca-1^+^ and PDGFRa^+^/Sca-1^-^ cells, which we found to be in close proximity (Fig. 1F and Suppl. Fig. S1F).

Thus, in the resting heart, both Sca-1^+^ and Sca-1^-^ cell types express similar transcriptional programmes indicative of a fibrogenic potential, and their transcriptome or anatomical location fails to point towards obvious functional differences between them.

Next, we assessed the clonogenic and developmental potential of PDGFRa^+^/Sca-1^+^ and Sca-1^-^ cells *in vitro*. Single tdTomato^+^ cells were seeded in 96-well plates containing C3H10T1/2 cells as a feeder layer and cultured for 14 days. Out of 288 wells, one in three Sca-1^+^ cells formed colonies and 25% of these colonies showed multi-lineage differentiation into both myofibroblasts and adipocytes (Fig. 1G). However, similar to what was reported by Chong *et al.*(Ieronimakis et al., 2013), we were unable to achieve reliable cell growth of Sca-1^-^ cells *in vitro* in clonal or bulk conditions, suggesting that this subset may be further differentiated (Suppl. Fig. S1G). To assess whether the PDGFRa, Sca-1^+^ cells contain all adipogenic activity detectable in the heart, bulk cultures from PDGFRaCre^ERT2^/Tdtomato mice, characterized in Suppl. Fig. 2A-B, were expanded in adipogenic media. As shown in figure 1H, the vast majority of perilipin^+^ adipocytes were also tdTomato^+^, suggesting that PDGFRa^+^ cells are the main contributor to adipocyte formation in the heart. Thus, a significant proportion of PDGFRa^+^/Sca-1^+^ cells, a population that has been previously defined as cardiac fibroblasts(Pinto et al., 2016), behave as multipotent mesenchymal progenitors *in vitro*, similar to skeletal muscle-derived FAPs(Joe et al., 2010; Uezumi et al., 2010).

**Figure 2.**
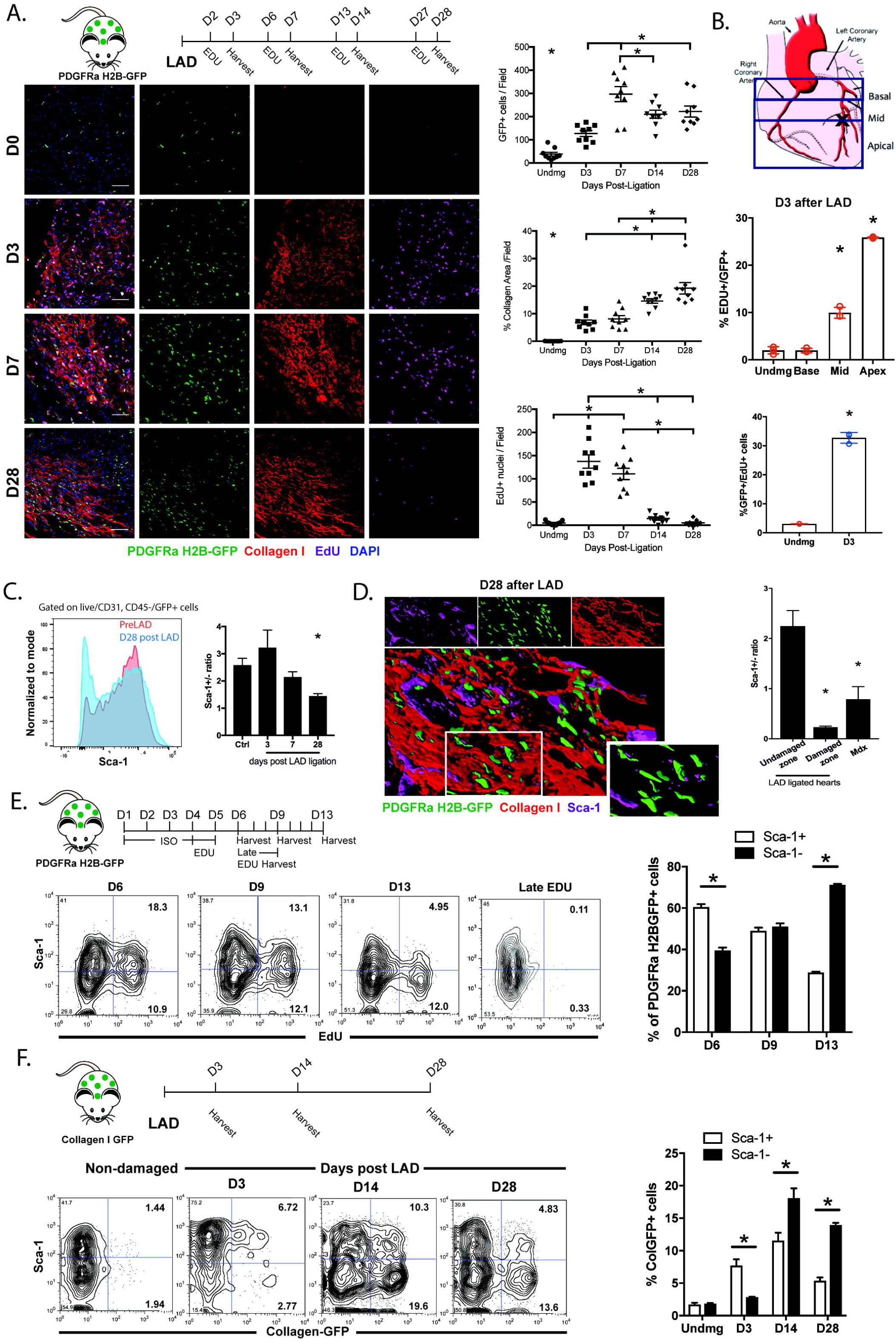
Response of cardiac PDGFRa+ cells to myocardial damage. A. Schematic of the experimental design for LAD ligation injury experiment in PDGFRa H2BGFP mice (top left). Representative images (bottom left) and quantification (right) of GFP^+^ cells, collagen I (red) and EdU (cyan) in 10µm cryosections from undamaged or LAD-ligated hearts at different time points post injury (n=9 per group; scale bar=60μm). B. Diagram to show where hearts, post LAD ligation, were cut and divided into apical, mid and basal sections (top). Quantification of percentage of PDGFRa H2BGFP^+^ that were also EdU^+^ in the apical, mid and basal sections of the heart 3 days post LAD ligation (mid; n=2 in each group). Quantification of the percentage of EdU^+^ cells that were also PDGFRa H2BGFP^+^ in the apical region of the heart 3 days post LAD ligation (bottom; n=3 in each group). C. Representative FACS histogram showing Sca-1 staining in undamaged (PreLAD) and 28 days LAD-ligated (D28 post LAD) hearts (left) and quantification of the Sca-1^+^/Sca-1^-^ cell ratio in the PDGFRa H2BGFP^+^ population (right). Only the apical region was used (n=3 in all groups except 28 days post LAD where n=4). D. Representative 3D reconstruction of a z-stack of images of collagen I (red) and Sca-1 (magenta) immunostaining of 10µm cryosections from PDGFRa H2BGFP mice hearts 28 days post LAD ligation (left). Inset: increased magnification of the area highlighted by the white rectangle showing PDGFRa H2BGFP and Sca-1 staining only. Scale bar = 20µm. Quantification of the Sca-1^+^/Sca-1^-^ cell ratio in the PDGFRa H2BGFP^+^ population in immunostained sections from 28 days LAD-ligated or 1 year old mdx mice hearts (right; n=3 for LAD-damaged and mdx hearts and n=4 for undamaged hearts). E. Schematic of the experimental design (top left). Representative FACS plots (bottom left) and quantification (right) of percentage of PDGFRa H2BGFP^+^ Sca-1^+^ or Sca-1^-^ that are EdU^+^ (relative to the total EdU+ cells) at different time points after ISO treatment for 5 days. D6, D9 and D13 are the 1^st^, 4^th^ and 8^th^ days, respectively, after the end of ISO treatment as shown in the schematic. F. Schematic of the experimental design (top left). Representative FACS plots (bottom left) and quantification (right) of percentage of Col1a1-3.6GFP^+^ that were Sca-1^+^ or Sca-1^-^ in undamaged hearts and at days 3, 14 and 28 post LAD ligation (n=4 in each group). Data represented as mean ± SEM, *p<0.05 compared to control undamaged group unless otherwise indicated by a horizontal bar.

### Stromal dynamics in the acutely damaged myocardium

To assess the response of cardiac PDGFRa^+^ cells to myocardial injury, we induced an experimental infarction by ligation of the left anterior descending coronary artery (LAD) in PDGFRa H2BGFP mice(Hamilton et al., 2003), characterized in Suppl. Fig. 2C. This resulted in a significant increase in PDGFRa cells (Fig. 2A), correlating with proliferation in the damaged region of the heart proximal to the apex, where more than 30% of EdU^+^ cells were GFP^+^ (Fig. 2A, B). In contrast to skeletal muscle injury, where upon successful regeneration FAPs numbers rapidly return to steady state levels, the increase in cFAPs after damage is sustained in time (Fig. 2A). Additionally, in contrast to undamaged areas where the majority of PDGFRa^+^ cells remained positive for Sca-1, we observed a time-dependent decrease in the ratio of Sca-1^+^ to Sca-1^-^ cells in the damaged heart apex (Fig. 2C). Since infarction often does not affect the full thickness of the left ventricular wall, we focused our analysis on the damaged areas, identified by the presence of collagen deposits. In these locations, more than 80% of the GFP^+^ cells failed to stain for Sca-1 (Fig. 2D and Suppl. Fig. S3A). Thus, Sca-1 negative cells increase upon damage, and they preferentially localize to areas of collagen deposition.

PDGFRa^+^ cells proliferate in response to cardiac damage. In order to more closely assess their proliferation kinetics, we switched to a model of damage amenable to higher throughput analysis. We injected PDGFRa H2BGFP mice with isoproterenol (ISO) for 5 consecutive days (100mg/kg, s.c.), a treatment known to induce subendocardial ischemic damage(Heather et al., 2009). EdU was administered (1mg/day) in the final 48 hours of ISO treatment. Twenty-four hours later, more than 50% of the EdU^+^ cells were GFP^+^ (Suppl. Fig. S3B). Whereas at this time point EdU incorporation was predominantly detected in Sca-1^+^ cells, seven days later most labeled cells were Sca-1^-^ (Fig. 2E). Importantly, no incorporation was observed when EdU was injected 3 days following the end of ISO treatment (Fig. 2E), indicating that any positivity observed past this time must be due to inheritance of the label from ancestry cells that were proliferating at the time of EdU administration. This strongly suggests that Sca-1^+^ cells preferentially respond to damage by proliferating, and that following initial expansion, they generate Sca-1^-^ progeny. The fact that Sca-1^-^ cells retain the label for over a week past the end of EdU administration additionally indicates that their proliferative capacity is limited, in agreement with our *in vitro* data.

The association of Sca-1^-^ cells with collagen-rich regions prompted us to investigate collagen expression in PDGFRa^+^ cells. To this end we induced myocardial damage in mice carrying EGFP under the control of a collagen I regulatory region activated in fibrogenic cells(Kalajzic et al., 2002). Similar to what was observed with EdU, early after damage EGFP expression was mainly associated with the Sca-1^+^ fraction, while at later stages it gradually shifted to Sca-1^-^ cells (Fig. 2F). This is consistent with Sca-1^-^ cells being the differentiated fibrogenic progeny of Sca-1^+^ progenitors.

TGFβ is known to induce a fibrogenic program in mesenchymal progenitors. To test whether this is associated with downregulation of Sca-1, we sorted PDGFRa^+^/Sca-1^+^ cells and placed them in culture. A small proportion of Sca-1^-^ cells were spontaneously generated during the first week of culture in the absence of TGFβ, but they did not increase over the course of the next two weeks (data not shown). As Sca-1^-^ cells do not expand efficiently in culture, this suggests that their numbers reach an equilibrium as a result of their continued production and dilution by proliferating Sca-1^+^ progenitors. Treatment with TGFβ for 7 days led to a doubling of the frequency of Sca-1^-^ cells in the cultures, consistent with the notion that Sca-1 downregulation parallels the adoption of a fibrogenic fate (Fig. 3A). In aggregate, this body of *in vitro* results supports the hypothesis that Sca-1^+^ cells represent progenitors for fibrogenic Sca-1^-^ cells.

**Figure 3.**
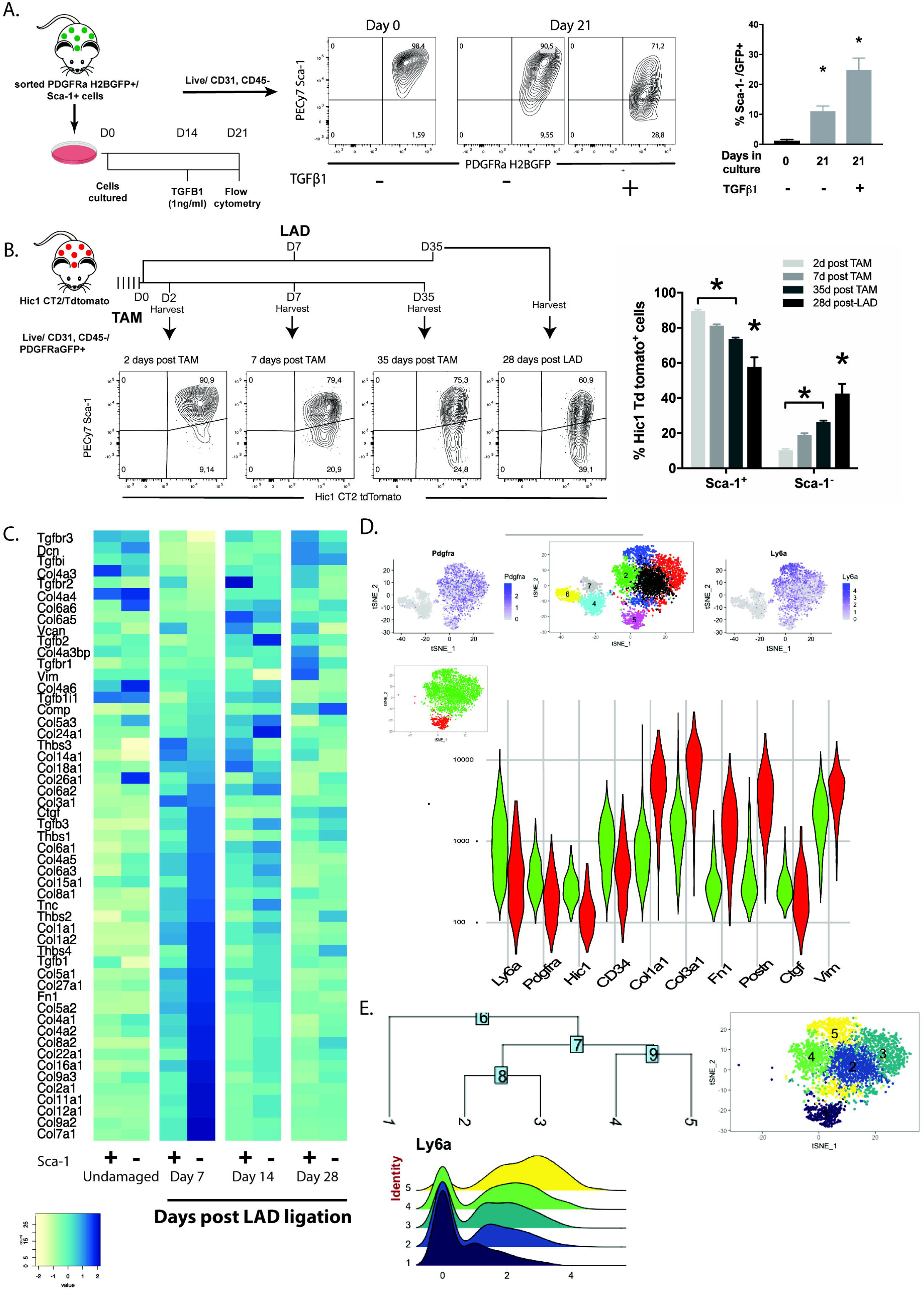
Sca-1+ cells represent progenitors for the more fibrogenic Sca-1-cells. A. Schematic of the experimental design (left). Representative FACS plots (middle) and quantification (right) of the percentage of PDGFRa H2BGFP^+^ cells that became Sca-1^-^ at day 21 in culture in the absence or presence of TGFß (1ng/ml) treatment for the last 7 days (n=3 in each group). B. Schematic of the experimental design (top left). Representative FACS plots (bottom left) and quantification (right) of the percentage PDGFRa H2BGFP^+^/Hic1Cre^ERT2^/tdTomato^+^ cells that were either Sca-1^+^ or Sca-1^-^ at different time points after TAM injection in undamaged or LAD ligated mice hearts (n=3 for 7days post TAM group and n=4 in all other groups). C. Heatmap of relative expression of key fibrogenic genes produced from sorted and RNA sequenced PDGFRa H2BGFP^+^ Sca-1^+^ or Sca-1^-^ cells in undamaged or LAD ligated hearts at different time points post injury. The heatmap was generated from the Z-score of the different genes at the specified time points (n=2 for undamaged and day 14 post LAD, n=3 for 7 days post LAD and n=4 for 28 days post LAD). D. t-SNE and violin plots of single cell RNA sequenced Hic1Cre^ERT2^/tdTomato^+^ cells isolated from the apical (damaged) region of the heart 7 days post LAD ligation. Top panel illustrates the expression of pdgfra and ly6a (Sca-1), while the bottom panel shows a violin plot of the expression of genes of interest in the 2 clusters expressing pdgfra. Data in the graphs are represented as mean ± SEM, *p<0.05 compared to all other groups or as indicated by a horizontal bar. E. cFAPs were subset from the Hic1Cre^ERT2^/tdTomato^+^ cells dataset shown in (D) and sub-clustered as shown in the t-SNE plot on the top right. A phylogenetic tree was constructed based on cluster similarity (left) and a ridge plot of the Ly6a (Sca-1) in the different cFAP clusters of the phylogenetic tree was drawn (bottom right).

We next tested this hypothesis in the context of cardiac damage, by means of lineage tracing in the Hic1Cre^ERT2^ mouse strain.. As expected, when animals were harvested immediately after the end of TAM treatment, tdTomato^+^ cells were almost entirely Sca-1^+^(D2 in Fig. 3B) confirming our findings with the Hic1 citrine GFP strain (Fig. 1C). Increasing frequencies of Sca-1^-^ cells were detected 7 and 35 days later ^+^(D7 and D35 in Fig. 3B), suggesting that, similar to what was observed *in vitro*, homeostatic production of these cells takes place in the absence of overt damage. LAD ligation significantly increased the generation of Sca-1^-^ cells, which reached approximately 50% of the labelled population 4 weeks after the surgery (Fig. 3B). A caveat to this experiment is that a small number of Sca-1^-^ cells is also labelled by Hic1Cre^ERT2^ immediately after TAM treatment, and therefore the observed results could also be explained by their preferential expansion following damage. To further investigate this possibility, we delivered 6-hour pulses of EdU to undamaged mice and found that the Sca-1^+^ showed significantly greater percentage EdU positivity (1.89±0.2%) compared to the Sca-1^-^ population (0.96±0.1%), ruling out the possibility that preferential proliferation was the cause of a shift in their ratio. Thus, both in homeostasis and at an increased rate after damage, Sca-1^+^ cFAPs generate Sca-1^-^ cells.

To examine in more depth the phenotype of the two cell types, we performed RNAseq on FACS-sorted PDGFRa H2BGFP^+^ cells that were either Hic1 tdTom^+^ Sca-1^+^ or Hic1 tdTom^-^ Sca-1^-^ before damage and at 1, 2 or 4 weeks post-LAD ligation and focused on the analysis of ECM-related genes. No obvious difference in gene expression was noticed between Sca-1^+^ and Sca-1^-^ at homeostasis or four weeks after surgery (Fig. 3C). However, expression of genes associated with pathogenic matrix deposition was significantly enriched in the Sca-1^-^ fraction 1 and 2 weeks following damage, confirming that these cells assume an activated fibrogenic phenotype (Fig. 3C). In accordance with this, t-SNE clustering of scRNAseq data from cardiac Hic1Cre^ERT2^/tdTomato^+^ cells 7 days after LAD ligation revealed 2 spatially distinct clusters expressing PDGFRa, one of which expressed lower levels of Sca-1 and Hic1 but was highly enriched for ECM-related genes as well as the activated fibroblast marker periostin(Kanisicak et al., 2016) (Fig. 3D).

Clustering analysis revealed that the PDGFRa+ cells retained their heterogeneity after damage (Fig. 3D) and that the cells expressing highest levels of Sca-1 were still clustered together (subcluster 3 shown in Fig. 3D and Suppl. Fig. S4A). Suppl. Fig. S4B shows a heatmap of the top 10 most differentially expressed genes in each of the PDGFRa^+^ subclusters.

Interestingly, when the cFAPs were sub-clustered and clusters were organized in a phylogenetic tree, the resulting hierarchy correlated with the levels of Sca-1 expression in each cluster (Fig. 3E) further supporting the hypothesis that cells downregulate Sca-1 as they differentiate along the fibrogenic lineage and suggesting that multiple intermediates may be generated during such transition.

Clustering of the merged scRNAseq datasets for Hic1Cre^ERT2^/tdTomato^+^ cells before and after damage (Suppl. Fig. S4B) revealed that FAPs from damaged samples clustered separately from those at steady state (Suppl. Fig. S4C). Gene Ontology Analysis of the differentially upregulated genes in each cluster (Suppl. Fig. S4D) pointed to the upregulation of multiple biosynthetic pathways upon FAP activation, and of ECM production-related as well as pro-angiogenic pathways upon loss of Sca-1 and differentiation.

Taken together, these results strongly suggest that in response to damage, cFAPs differentiate into fibrogenic cells while downregulating progenitor markers such as Sca-1 and Hic1.

### Inhibition of FAPs differentiation improves cardiac performance

Excessive generation of fibrogenic matrix causes heart wall stiffening, resulting in impaired cardiac contractile function(Gyöngyösi et al., 2017). We reasoned that inhibiting the differentiation of cFAPs towards Sca-1^-^ fibrogenic cells may have a therapeutic effect following myocardial infarction. We previously identified members of the imatinib/nilotinib tyrosine kinase inhibitor (TKI) family as modulators of FAP survival(Lemos et al., 2015). Further analysis revealed that these compounds are also capable of blocking the upregulation of collagen in TGFβ-treated cFAPs *in vitro* (Fig. 4A). Thus, we investigated whether nilotinib (25mg/kg/day) could ameliorate ISO-induced cardiac damage. Mice were treated with ISO for 5 days, and with nilotinib for the following 7 days. The latter treatment prevented the decrease in the Sca-1^+^/Sca-1^-^ ratio observed with ISO alone (Fig. 4B) and significantly reduced fibrotic lesion formation (Fig. 4C, D), supporting the hypothesis that nilotinib acts, at least in part, by blocking pro-fibrotic differentiation of cFAPs and loss of Sca-1 expression. As cardiac function was not significantly impacted by ISO treatment, we sought to establish the therapeutic potential of blocking cFAP-driven fibrogenesis in the more severe LAD coronary artery ligation model. Three days after surgery, mice were treated with nilotinib (25mg/kg/day) for 28 days, after which cardiac function was assessed. We delayed the start of the treatment to avoid interfering with the initial steps of scar formation, as this has been shown to be deleterious(Frantz et al., 2008; IKEUCHI et al., 2004), and to more closely mimic the likely timing of administration of similar drugs to human patients.

**Figure 4.**
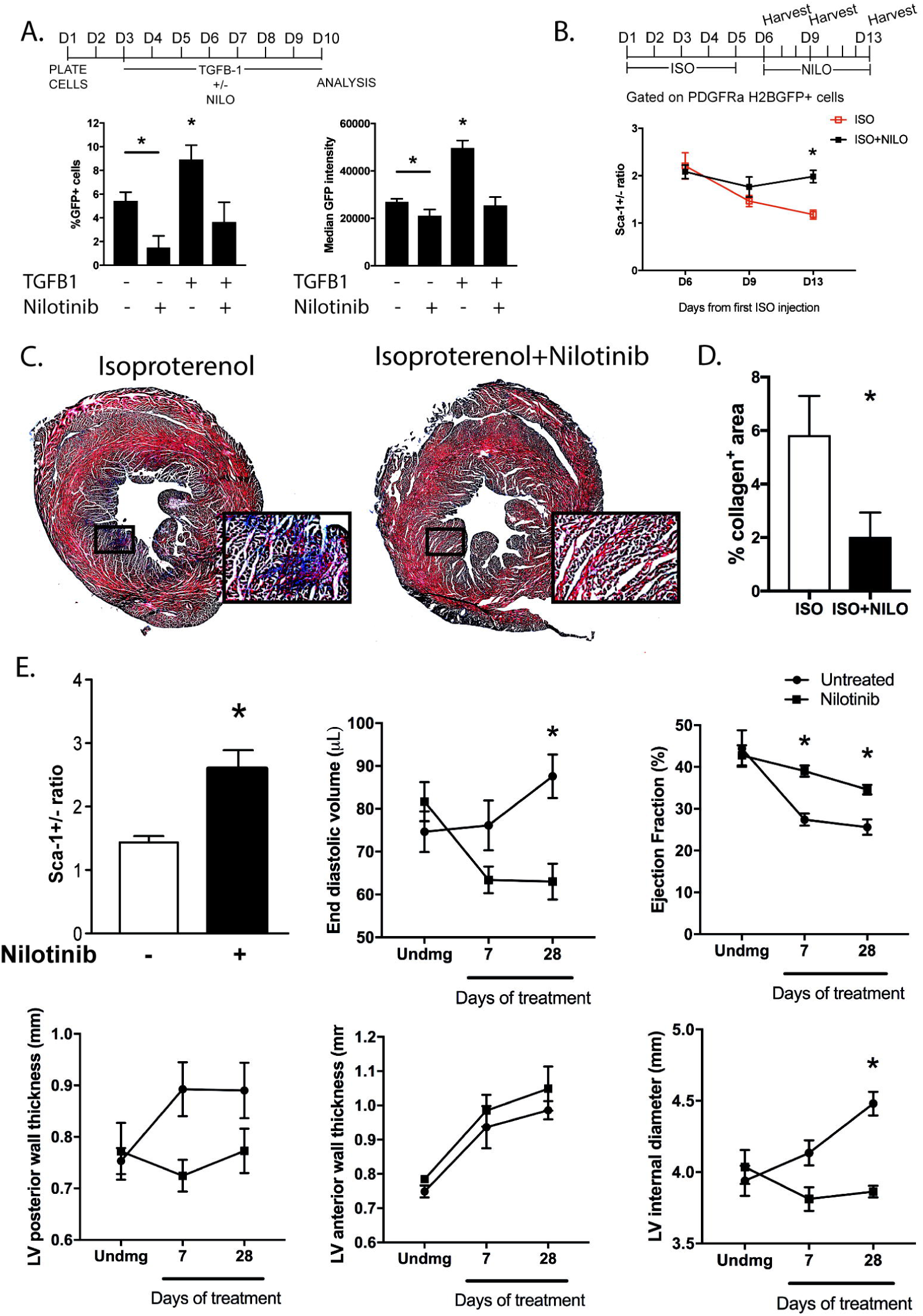
Inhibition of FAPs differentiation improves cardiac performance. A. Schematic of the experimental design and quantification of the percentage of Col1a1-3.6GFP^+^ cells and median GFP intensity in sorted Sca-1^+^ cells cultured for 7 days in the absence or presence of TGFß (1ng/ml) +/-nilotinib (1µM; n= 6 wells for control and TGF groups and n=7 in nilotinib only and n=9 in TGF+nilotinib). B. Schematic of experimental design and quantification of the Sca-1^+^/Sca-1^-^ cell ratio within the PDGFRa H2BGFP^+^ population in the heart at different time points after ISO treatment (n=3 in all groups except D13 groups where n=4). C. Photomicrographs of transverse sections of hearts from ISO-treated mice +/-nilotinib stained with Masson’s trichrome staining. The insets are zoomed in views of the areas outlined by the black rectangles. D. Representative images and quantification of the area of collagen I immunostaining in heart cryosections of PDGFRa H2BGFP mice treated with ISO (100mg/kg) +/-nilotinib (25mg/kg; n=5 in each group). E. Quantification of the effect of nilotinib treatment on the Sca-1^+^/Sca-1^-^ ratio within the PDGFRa H2BGFP^+^ population and on cardiac function post LAD ligation. Cardiac function was assessed by echocardiography and the end diastolic volume, ejection fraction and left ventricular chamber dimensions were measured (n=5 for undamaged, 6 for 7 days untreated, 7 for 7 and 28 days nilotinib-treated and 11 for 28 days untreated group). Data are represented as mean ± SEM. *p<0.05 compared to untreated group (at same time point in the line graphs).

In the untreated mice, LAD ligation produced a progressive decline in LV ejection fraction. This was associated with elevated end-diastolic volume (a hallmark of cardiac compromise), as well as increased anterior and posterior LV wall thickness and LV chamber dilation, features of cardiac remodeling (Fig. 4E). Interestingly, nilotinib treatment attenuated the decline in ejection fraction and completely prevented the elevation in end-diastolic volume, LV chamber dilation and LV posterior wall thickening. Similar to ISO injury model, nilotinib treatment was also associated with a significant increase in the PDGFRa^+^ Sca-1^+^/Sca-1^-^ ratio (Fig. 4E). Thus, the differentiation step leading to the generation of fibrogenic cells from cFAPs can be pharmacologically targeted to improve post-MI cardiac performance.

### A role for cFAPs in the pathogenesis of arrhythmogenic cardiomyopathy (AC)

While adipogenic differentiation of cFAPs was readily observed *in vitro*, their activation following ischemic damage does not lead to the appearance of intramyocardial adipocytes. However, adipocyte accumulation is a well-documented outcome of Arrhythmogenic Cardiomyopathy (AC), an inherited cardiac disease characterized by degenerative replacement of myocardium with fibrofatty infiltrations leading to life threatening arrhythmias and sudden death, especially in young adults(Basso et al., 2009).

To confirm the adipogenic potential of cFAPs *in vivo*, we generated a model of this disease by overexpressing human desmoglein 2 (DSG2) carrying a pathogenic 1672C →T mutation(Pilichou et al., 2006) in mice. In this model, fibrous and/or fatty infiltrations develop by 9 months of age, and adipocytes have been shown to ail from cells originating from a PDGFRa precursor. However, as a non-inducible PDGFRa-CRE was used, it is unclear if this marker expression took place during development or in the adult heart(Lombardi et al., 2016). To assess whether such infiltrations originate from Hic1^+^ cells, we bred the tdTomato reporter and either the Hic1Cre^ERT2^ or the PDGFRaCre^ERT2^ transgenes into this strain (Fig. 5A, D). Consistent with the ischemic injury models, Hic1 or PDGFRa traced cells were enriched in areas of collagen deposition (Fig. 5B, E). In addition, the majority of infiltrating adipocytes were tdTomato^+^ (Fig. 5C, F) confirming that PDGFRa^+^ Hic1^+^ progenitors are the origin of fibrofatty depositions in AC, and that cFAPs are indeed adipogenic *in vivo*.

**Figure 5.**
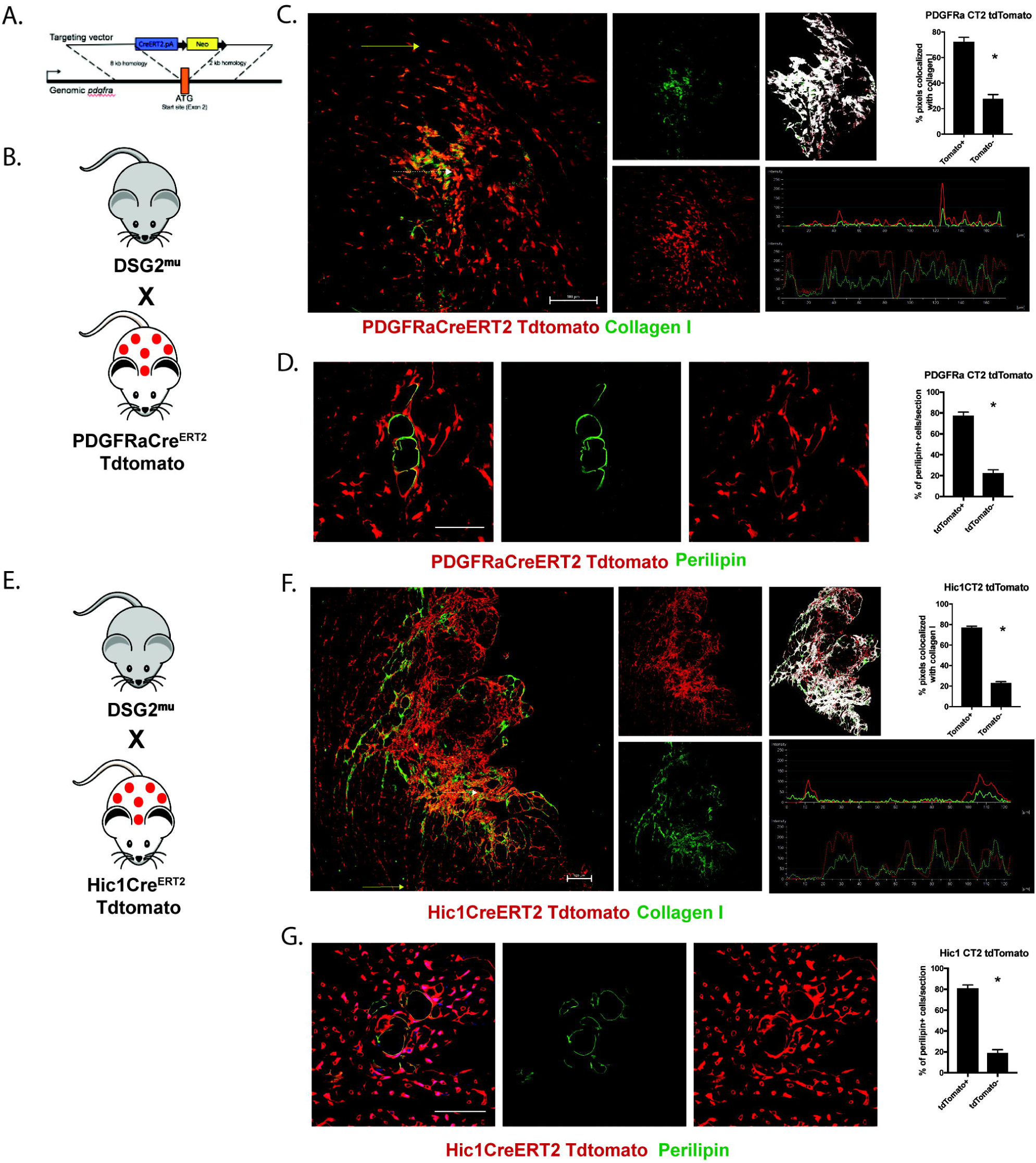
Lineage tracing of fibrous and fatty infiltrations in a murine model of arrhythmogenic cardiomyopathy (AC) using PDGFRa or Hic1 Cre systems. A. Schematic of the PDGFRa Cre^ERT2^ “knock-in” allele. Breeding scheme for generation of DSG2^mu^/PDGFRaCre^ERT2^/tdTomato (B) or DSG2^mu^/Hic1Cre^ERT2^/tdTomato (E) mice for lineage tracing. Representative images and quantification of the PDGFRa/tdTomato^+^ or tdTomato^-^ (C) or Hic1/tdTomato^+^ or tdTomato^-^ cells (F) that were colocalized with collagen I in the fibrous lesions in DSG2^mu^ mice hearts. Colocalization was measured using imageJ ^®^and a representative colocalized pixel map image generated by the software in the fibrous lesion is shown. The line graphs in (C) and (F) represent the intensity profile of tdTomato (red) and collagen I (green) staining across the arrows shown in the merged images; the solid line graphs are for the solid arrow (representing an area of no damage) and the broken line graphs are for the broken arrow (an area of damage; n=4 in each group). Representative images and quantification of the PDGFRa/tdTomato^+^ or tdTomato^-^ (D) or Hic1/tdTomato^+^ or tdTomato^-^ cells (G) that were also perilipin^+^ in the adipocytic infiltrations in DSG2^mu^ mice hearts (n=4 per group for Hic1 traced cells and n=6 per group for PDGFRa traced cells); scale bar= 60 μm. Data are represented as mean ± SEM. *p<0.05 compared to respective tdTomato^-^ group.

### Hic1 deletion leads to activation of cFAPs and development of AC pathology in the absence of cardiomyocyte damage

The lack of adipocytic infiltrations in ischemic damage and their abundance in mice carrying the pathogenic DSG2 mutation suggest that the differentiated output of cFAPs is influenced by the type of injury. Next, we sought to assess their developmental potential in the absence of any overt (LAD ligation) or subclinical (AC) cardiomyocyte damage. We have recently found that Hic1 controls quiescence of mesenchymal progenitors (Scott *et al.*, in preparation) and that its deletion leads to spontaneous FAP activation in multiple tissues. Indeed, in UBCCre^ERT2^ Hic1^Fl/LacZ^ mice, Hic1 deletion resulted in doubling of Xgal^+^ cells as early as 10 days after TAM induction (Fig. 6A). Immunohistochemical analysis of Hic1 KO, but not control heart sections 6 months to 1 year after TAM treatment revealed epicardial and/or interstitial fibrosis as well as intramyocardial adipocyte infiltrations (Fig. 6B). In these mice, cardiac function was compromised as illustrated by reduced LV Ejection fraction as well as increased myocardial performance index (MPI), isovolumic relaxation time (IVRT) and mitral valve (MV) deceleration time (Fig. 6C, D). Additionally, ECG tracings of Hic1 KO mice hearts showed arrhythmias in the form of premature ventricular contractions as well as notched QRS complexes (white arrows in Fig. 6C). To evaluate the systemic effects of these changes, we subjected these mice to forced exercise and assessed their endurance. The distance and speed reached until fatigue (as defined in the methods section) were significantly lower in Hic1 KO mice than in the control mice (Fig. 6E).

**Figure 6.**
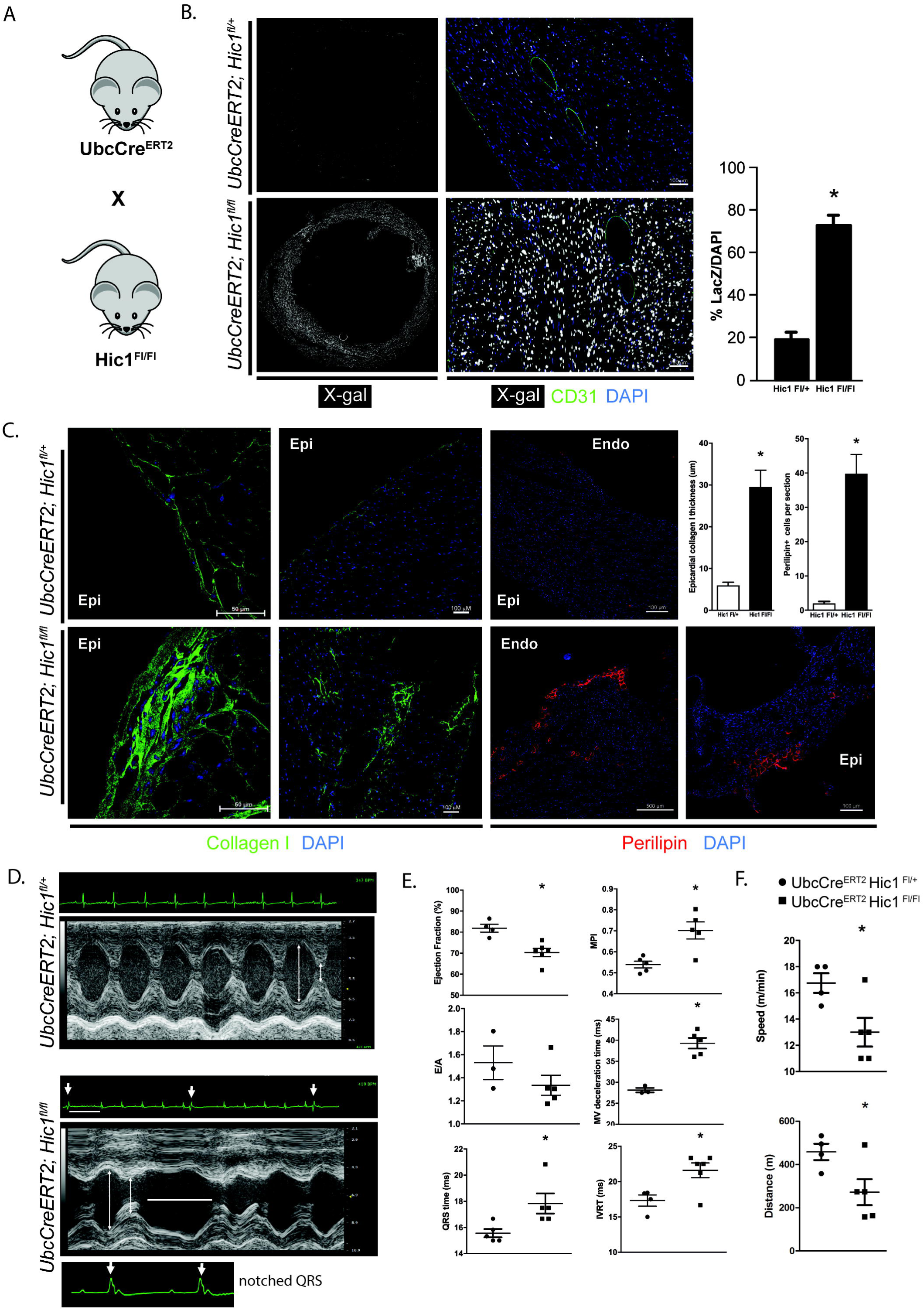
Hic1 deletion leads to activation of cFAPs and AC phenotype in the absence of cardiomyocyte damage. A. Breeding scheme for generating UBCCre^ERT2^/Hic1^Fl/Fl^ mice harboring also LacZ gene under the control of the Hic1 promoter. B. Representative images and quantification of the LacZ^+^ cells in UBCCre^ERT2^/Hic1^Fl/Fl^ mice hearts and their controls 10 days after TAM-induced recombination (n=5 per group). C. Representative images showing epicardial and interstitial fibrosis, and perilipin^+^ adipocyte infiltrations in Hic1-deleted mice hearts 1 year after TAM-induced recombination (Epi=epicardial side, Endo=endocardial side) and quantification of the epicardial layer thickness and perilipin^+^ cells in these mice hearts and littermate controls (n=3 in the Hic1^fl/+^ and n=5 in Hic1^fl/fl^ group). D. Representative M-mode echocardiographic and ECG tracings of UBCCre^ERT2^/Hic1^Fl/Fl^ mice hearts and their controls. White arrows and horizontal lines highlight the premature ventricular extra beats and associated refractory pause, respectively. The white arrowheads highlight the notched QRS complexes found in the Hic1 knockout mice. E. Quantification of cardiac function in Hic1 knockout mice and their controls. Echo was done and ejection fraction, myocardial performance index (MPI), E/A ratio, isovolumic relaxation time (IVRT) and mitral valve (MV) deceleration time were calculated to determine systolic and diastolic function. QRS time was calculated from the ECG tracings performed during the echo procedure (n=4 for Hic1^fl/+^ group and n=6 for Hic1^fl/fl^ group). F. Quantification of the speed and distance reached by mice force-exercised on a treadmill as indices of physical endurance in Hic1 knockout mice and their controls (n=4 for Hic1^fl/+^ and n=5 for Hic1^fl/fl^ group). Data are represented as mean ± SEM. *p<0.05 compared to UBCCre^ERT2^/Hic1^Fl/+^ control mice.

The UBCCre^ERT2^ Hic1^Fl/LacZ^ deletes Hic1 in both PDGFRa positive and negative cells, making it more difficult to attribute the observed phenotype exclusively to cFAPs versus the minor, Hic1 expressing pericytic population described above. To bypass this confound, we bred Hic1^Fl/Fl^ mice with PDGFRaCre^ERT2^/tdTomato mice and confirmed that the vast majority of cells composing the lesions originated from PDGFRa^+^ cFAPs (Suppl. Fig. S5A). As expected, these mice also developed signs of cardiac dysfunction comparable with those described above (Suppl. Fig. S5B and C). To further explore whether Hic1 may be a modifier of the penetrance of AC, we crossed the DSG2^mu^ mice with PDGFRaCre^ERT2^/Hic1^Fl/Fl^ mice and compared them to their parental strains. At 6 months of age and 1 month after TAM treatment, adipocytic infiltration was limited in the DSG2^mu^ mice (as at this age DSG2^mu^ mice do not display overt lesions yet), while significantly increased in the other two strains, with mice harboring both the DSG mutation and the Hic1 KO having approximately double the adipocytes than the Hic1KO mice (Suppl. Fig. S5D). Thus, damage-independent activation of FAPs leads to fibrous/fatty infiltration and arrhythmias reminiscent of human AC even in the absence of cardiac damage, and accelerates fat deposition in the presence of subclinical damage, underpinning the key role played by these cells in AC pathophysiology.

## Discussion

The cellular mechanisms leading to cardiac fibrosis and fatty degeneration remain poorly understood. Over the past several years, various markers have been proposed to identify cardiac fibroblasts, including multiple lineage tracing tools, such as PDGFRa, TCF21 and collagen1. However, despite reports that multipotent progenitors are also contained within these populations(Chong et al., 2011), their phenotypic heterogeneity and functional properties have not been explored in depth.

Here we show that a significant portion (∼25%) of the PDGFRa^+^ cells capable of forming colonies *in vitro* represent mesenchymal progenitors with fibroadipogenic potential. We further show that such potential is not limited to *in vitro* conditions but is expressed *in vivo*. Based on these findings, we suggest the nomenclature be changed to reflect the fact that, like in skeletal muscle, the Sca-1^+^ subset of cardiac fibroblasts are in fact multipotent mesenchymal progenitors. In skeletal muscle, FAPs can also respond to exogenous BMPs or pathogenic alterations to generate ectopic bone(Wosczyna et al., 2012). While we have not investigated the osteogenic potential of their cardiac counterparts at the clonal level, Hic1^+^ progenitors are involved in developmental osteogenesis as well as bone regeneration in the adult (Scott *et al*. in preparation), suggesting that the recently reported osteogenic potential of cardiac fibroblasts(Pillai et al., 2017) may at least in part reside within this population. Our lineage tracing experiments exclude any significant contribution of PDGFRa^+^ or Hic1^+^ cells to cardiomyocytes or endothelial cells, either spontaneously or following different types of damage. Thus, unlike what was previously described for mesenchymal stem cells(Noseda et al., 2015; Ubil et al., 2014), their multipotency is restricted to connective tissue elements. In view of their predominant fates *in vitro* and *in vivo*, and in keeping with the nomenclature in use in skeletal muscle, we propose that PDGFRa, Sca-1, Hic1 positive cells be called cardiac fibro/adipogenic progenitors (cFAPs) as used in this study.

We identify a differentiation hierarchy within cardiac PDGFRa^+^ cells, with cFAPs constantly generating a small number of post-mitotic, Sca-1^-^ differentiated cells in the undamaged heart. A caveat of this interpretation lies in the fact that TAM treatment, at the dosage used, can be cardiotoxic(Bersell et al., 2013). At homeostasis, both cFAPS and their progeny express low levels of pathogenic ECM genes, but following damage these genes, as well as the recently proposed activated fibroblast marker periostin(Kanisicak et al., 2016), are highly upregulated in the differentiated Sca-1^-^ cells. Over the process of damage repair, such fibrogenic transcriptional programme is gradually downregulated to the point where little difference remains between Sca-1^+^ progenitors and Sca-1^-^ progeny after 1 month of damage. This validates the proposed existence of cardiac fibrogenic cells in at least two states, deactivated and active, and it suggests that therapeutic interventions modulating fibrosis should be deployed not immediately after an ischemic event to allow for scar formation, but rather within the first few days following damage.

Inhibition of such differentiation step using nilotinib, a second generation anti-leukemic TKI, led to an improvement of heart function following an ischemic infarct and reduced the generation of Sca-1^-^ cells from cFAPs. Consistent with these results Imatinib, an earlier member of the same TKI family, was shown to also have favorable effects on rat cardiac function after MI(Liu et al., 2014). While our experiments provide clear proof of principle that modulation of cFAP differentiation is a viable therapeutic strategy, the reported cardiotoxicity of nilotinib(Moslehi and Deininger, 2015) raises questions about whether its long term use in infarct patients is appropriate, and provides a strong rationale for testing alternative TKIs.

By performing lineage tracing in a mouse model of AC, we show that the Hic1^+^ subset of PDGFRa^+^ cells is responsible for the generation of the intramyocardial adipocytes pathognomonic of this disease. This is in stark contrast with these cells’ exclusive generation of fibrogenic progeny in the context of myocardial infarction, suggesting that the extent or the type of damage inflicted is likely to modulate their lineage choice through mechanisms as yet unclear. This is consistent with the upregulation of known anti-adipogenic factors such as TGFβ and Wnts following an MI(Blankesteijn et al., 2000; Deten et al., 2001).

cFAPs’ quiescence can be prevented by deleting Hic1, a transcriptional repressor that modulates the expression of cell-cycle related genes (Scott et al, in preparation). In the heart, this led to a rapid increase in cFAP numbers followed by the slower appearance of intramyocardial adipocytes and fibrosis, and eventually clinical signs reminiscent of human AC and reduced running performance. This suggests that the main effects of the desmosomal mutations linked to AC may be to cause the chronic activation of cFAPs, which represents the key pathogenic event. Consistent with this hypothesis, lack of Hic1 cooperates with AC-inducing mutations in accelerating the appearance of intramyocardial adipocytes.

Our work reveals that a significant proportion of cells meeting the phenotypic definition of cardiac fibroblasts are actually multipotent progenitors, identifies the quiescence regulator Hic1 as a marker for these cells, and points to their significant role in the pathology of human AC. It also suggests that following ischemic damage, TKIs can be used to modulate the appearance of fibrogenic cells with positive results on cardiac performance, providing a rationale for exploring a novel therapeutic strategy for one of the most common diseases of our time.

## Methods

### Animals

PDGFRa H2BGFP mice were purchased from Jackson Laboratories and Col1a1-3.6GFP mice were a gift from Dr. David W. Rowe (Center for Regenerative Medicine and Skeletal Development, University of Connecticut Health Center). For generation of PDGFRa Cre^ERT2^ “knock-in” allele, a Cre^ERT2^ polyA cassette and a FRT-flanked neo cassette were recombined into the start codon of Pdgfra. The construct was electroporated into TL1 (129S6/SvEvTac) ES cells and these were injected into C57BL/6 blastocysts. The neo cassette was removed. PDGFRa Cre^ERT2^ mice are maintained on a C57BL/6 background. Mice harboring Hic1 Cre^ERT2^, or Hic1^Fl/Fl^ alleles were generated in the Underhill lab while mice harboring Hic1 citrine were generated by the Korinek lab(Pospichalova et al., 2011). Hic1 Cre^ERT2^ mice were crossed with tdTomato mice and Hic1^Fl/Fl^ mice were crossed with UBC Cre^ERT2^ or PDGFRa Cre^ERT2^ mice. Transgenic mice with cardiac-specific overexpression of FLAG-tagged wild type and mutant human DSG2 (c.1672C>T, p.Q558X; DSG2^mu^) were generated. Full-length WT and mutant human DSG2 cDNAs were expressed under the control of mouse alpha-myosin heavy chain promoter. The transgenic mice were maintained on a C57BL/6 background. DSG2^mu^ mice were crossed with Hic1 Cre^ERT2^ or PDGFRa Cre^ERT2^ mice. Allelic recombination under the PDGFRa Cre^ERT2^, Hic1 Cre^ERT2^ or UBC Cre^ERT2^ alleles was induced by daily injections of 0.1mg/g TAM for 5 days, administered intraperitoneally. To control for TAM toxicity, all mice, including controls, were administered TAM. Cardiac damage was induced by subcutaneous injection of 100 mg/kg of ISO (Sigma) for five consecutive days. In vivo nilotinib (Novartis) treatment was achieved by daily intraperitoneal injections of 25 mg/kg at indicated times. To experimentally induce myocardial infarctions, mice were intubated, anesthetized and their left anterior descending artery ligated as previously described(Kolk et al., 2009). EdU (Life Technologies) was given for indicated time periods at 1 mg/mouse injected intraperitoneally. Experiments were performed in accordance with University of British Columbia Animal Care Committee approved protocols.

### Heart digestion and flow cytometry

Mice were sacrificed, their hearts carefully excised and cut into 2mm pieces. A single cell suspension was made by digesting tissue for 30 minutes at 37°C in Collagenase type 2 solution (Worthington Biochemical; 500uL per heart of 2.5 U/ml) in 10 mM CaCl2. Suspensions were then centrifuged at 800 rpm and supernatant decanted. This was repeated twice followed by incubation for 1 hour at 37°C in a solution containing Collagenase D (Roche; 1.5 U/ml), Dispase II (Sigma-Aldrich; 2.4 U/ml) and 10 mM CaCl_2_. Following washing and filtration with 70 µm mesh filters, cell preparations were incubated with primary antibodies against cell membrane markers for 30 min at 4 °C in supplemented PBS containing 2 mM EDTA and 2% FBS (FACS buffer) at ∼3 × 107 cells/ml. The antibodies used in flow cytometry and the dilutions are listed on Table S1 of Supplementary Information.

For in vivo proliferation assays, EdU (Thermo Fisher Scientific) was dissolved in PBS at 2 mg/ml solution and it was administered daily by intraperitoneal injection (40 mg/kg). For flow cytometric analysis, cells were first stained for surface markers, then stained for EdU incorporation using the Click-iT™ Plus EdU Pacific Blue™ Flow Cytometry Assay Kit as described in the manufacturer’s manual. Analysis was performed on LSRII (Becton Dickenson) equipped with three lasers. Data were collected using FacsDIVA software. Cell sorting was performed on a FACS Influx (Becton Dickenson) or FACS Aria (Becton Dickenson) sorters. Sorting gates were strictly defined based on fluorescence minus one stains. Flow cytometry data analysis was performed using FlowJo 10.4.1 (Treestar) software.

### Quantitiative Reverse Transcription PCR

RNA isolation was performed using RNeasy mini kits (Qiagen) and reverse transcription was performed using the Superscript Reverse Transcriptase (Applied Biosystems). Fibrogenic gene expression analysis was performed using Taqman Gene Expression Assays (Applied Biosystems), on a 7900HT Real Time PCR system (Applied Biosystems). Sequence information for the primers contained in the Taqman assays are not in the public domain, but ordering information is provided in Supplementary Information. Data were acquired and analyzed using SDS 2.0 and SDS RQ Manager software (Applied Biosystems).

### Histology and immunohistochemistry

Mice received an intraperitoneal injection of 0.5 mg/g Avertin, and were perfused transcardially with 20 ml PBS/ 2 mM EDTA, followed by 20 ml of 4% paraformaldehyde (PFA). The excised muscles were fixed in 4% PFA overnight. The following day, the tissue was either prepared for paraffin embedding following the standard sequence of dehydration steps in ethanol, or transferred to 30% sucrose in PBS and incubated overnight for subsequent optimal cutting temperature (OCT) formulation embedding and freezing. Standard methods were followed for cryosectioning and 10 µm tissue sections were either stained using Masson’s Trichrome Stain as previously described(Bostick et al., 2012), or processed for immunohistochemistry as follows: tissue sections were washed in 0.3% Triton X-100 (Sigma) in PBS, incubated in sodium borohydride solution (10 mg/ml) for 30 minutes and then blocked for 1 hour at room temperature in PBS containing 10% normal goat serum (NGS), 0.2% Triton X-100 and 5% bovine serum albumin (BSA). Sections were stained overnight at 4 °C using a primary antibody diluted in 10% NGS, 0.2% Triton X-100 and 5% BSA. The primary antibodies used for immunofluorescence are listed in Table S1 of Supplementary Information. In all cases, the primary antibody was detected using secondary antibodies conjugated to Alexa 488, 555, 594, or 647 (Molecular Probes). Confocal microscopy was performed using a Nikon eclipse Ti Confocal Microscope with a C2 laser unit. Figures were assembled using Illustrator CS6 (Adobe).

### Echocardiography

Cardiac function and dimensions were evaluated by two-dimensional transthoracic echocardiography. Echocardiographic image acquisition was carried out using the Vevo2100 system (Fujifilm Visualsonics). Mice were anaesthetized with 3-4% isoflurane in 100% oxygen. Following induction, mice were placed on a heated handling table and anesthesia and body temperature were maintained at 1-1.5% isoflurane and 37 ± 0.5°C, respectively. All measurements were performed by a single experienced operator blinded to the mouse genotypes.

### *In vitro* culturing experiments

Cells were sorted from digested hearts using indicated gating into high (4.5g/L) glucose Dulbecco’s modified eagle medium (DMEM, Thermo Fisher), supplemented with 10% (v/v) fetal bovine serum (FBS), 10 ng/ml bFGF (Peprotech), 1% (v/v) Penicillin Streptomycin (ThermoFisher) and 2 mM L-glutamine (Thermo Fisher). Media was changed every 2-3 days. For adipogenic differentiation, following 7 days of growth, medium was changed to MesenCult™ MSC basal medium (Stem Cell Technologies) supplemented with MesenCult™ adipogenic stimulatory supplement for mouse (Stem Cell Technologies) and 2 mM L-glutamine. Cells were cultured for an additional 14 days and adipogenesis was assessed via immunostaining for the adipocyte marker perilipin.

For *in vitro* differentiation studies, hearts from Col1a1-3.6GFP mice were digested and Col GFP-cells were grown for 3 days in high (4.5g/L) glucose DMEM (Thermo Fisher), supplemented with 10% v/v fetal bovine serum (FBS), 10 ng/ml bFGF (Peprotech) and 2 mM L-glutamine (Thermo Fisher) and treated with TGFß1 (1ng/ml), and/or nilotinib (1uM) for one week before being analyzed.

For clonal expansion and developmental potential experiments, FACS sorted single PDGFRaCre^ERT2^/ tdTomato^+^/Sca-1^+^ cells were seeded in 96 well plates on a feeder layer of gamma irradiated mouse-derived C3H/10T1/2 cells. The cells were cultured in high (4.5g/L) glucose DMEM (Thermo Fisher), supplemented with 20% v/v fetal bovine serum (FBS), 10 ng/ml bFGF (Peprotech), 2 mM L-glutamine (Thermo Fisher) and 1% (v/v) Penicillin Streptomycin (ThermoFisher) for 14 days. Afterwards, half of the cells were cultured in in high (4.5g/L) glucose DMEM (Thermo Fisher), supplemented with 5% v/v fetal bovine serum (FBS), 10 ng/ml bFGF (Peprotech) and 2 mM L-glutamine (Thermo Fisher) and the other half cultured in MesenCult™ MSC basal medium (Stem Cell Technologies) supplemented with MesenCult™ adipogenic stimulatory supplement for mouse (Stem Cell Technologies) and 2 mM L-glutamine for another 14 days. Colonies, defined as ≥ 50 red cells per well, were counted and, following fixation with 2% paraformaldehyde, were immunostained for perilipin (adipocytes) or aSMA (myofibroblasts).

### Quantification of tdTomato colocalization with collagen I

Colocalization with collagen I immunostaining in heart sections was determined using ImageJ (Fiji v1 edition). Split channel images were generated and the area of fibrosis was highlighted. Colocalization was determined using the colocalization thresholds plugin and a colocalized pixel map image was generated with constant intensity for colocalized pixels. % colocalization was determined as the % channel volume (number of voxels which have both channel 1 (collagen I) and channel 2 (tdTomato) intensities above threshold determined using Pearson’s coefficient, expressed as a percentage of the total number of voxels for collagen I channel above its threshold.

### Treadmill forced exercise test

Hic1 KO mice and their controls were subjected to low intensity short periods of exercise for 5 consecutive days for training purpose on a rodent treadmill equipped with shock grids to deliver very low level electrical stimulus when mice stop running (Columbus Instruments). After 2 days of rest, mice were run on the treadmill at a constant velocity of 10m/min and the distance travelled before fatigue (the time point at which mice are electrically shocked 3 consecutive times and do not respond by continuing to run) was determined. Alternatively, mice were subjected to increasing treadmill velocities (1m/min incremental increase every 50-meter run) and the velocity at which mice displayed fatigue was determined.

### *in situ* X-gal staining

For in situ X-gal staining, mice were terminally anesthetized by intraperitoneal injection of Avertin (400 mg/kg), the chest was opened and hearts were perfused with cold LacZ fixative (100 mM MgCl_2_, 0.2 % glutaraldehyde, 5 mM ethylene glycol tetra-acetic acid in PBS), then dissected and immersed in LacZ fixative for 3 hours on ice. For cryosectioning, samples were washed 3 times for 30 mins with PBS then incubated through a cryoprotective series of sucrose solutions of increasing concentration from 10-50% for ≥3 hours each before embedding into OCT compound (Tissue Tek) in disposable cryomolds (Polysciences) and frozen in an isopentane bath cooled by liquid nitrogen. Cryosections were cut at a thickness of 5-30μm and mounted onto Superfrost Plus slides (VWR). For detection of LacZ on sections, in situ LacZ staining with X-gal was carried out, where slides were incubated overnight at 37oC in a humidified chamber with the LacZ staining solution (2 mM MgCl_2_, 0.01 % Deoxycholate, 0.02% NP40, 5 mM potassium ferricyanide, 5 mM potassium ferrocyanide and 1mg/mL 5-bromo-4-chloro-3-indolyl-β-D-galactopyranoside). Slides were counter-stained with nuclear fast red and mounted with Aqua Polymount.

### Gene expression profiling – popRNA-seq

Total RNA was isolated using RNeasy Plus micro kit (Qiagen) as per manufacturer’s instructions. Sample integrity was tested on an Agilent Bioanalyzer 2100 RNA 6000 Nano kit (Agilent). RNA samples with a RNA Integrity Number > 8 were used for library preparation using the protocol for the TruSeq Stranded mRNA library kit (Illumina) on the Illumina Neoprep automated microfluidic library prep instrument. Paired end sequencing was performed on the Illumina NextSeq 500 using the High Output 150 cycle Kit (Illumina).

### RNA-seq Bioinformatic Analyses

De-multiplexed read sequences were then aligned to the Mouse Genome mm10 reference sequence using TopHat splice junction mapper with Bowtie 2 (http://ccb.jhu.edu/software/tophat/index.shtml) or STAR (https://www.ncbi.nlm.nih.gov/pubmed/23104886) aligners. Assembly and differential expression was estimated using Cufflinks (http://cole-trapnell-lab.github.io/cufflinks/). Heatmap generation were performed with VisRseq(Younesy et al., 2015) (v.0.9-2.15).

### Single Cell RNA-seq

Single cell suspensions were prepared by digesting the hearts and FACS-sorting target population into sterile-filtered culture medium (DMEM containing 5% FBS). Second sorting in 35μL medium was done to ensure high purity and quality of the cells. After the tdTomato^+^ cell suspension was examined for quality and cells counted using a hemocytometer, they were delivered into a Chromium Controller (10_X_ Genomics), captured and libraried with the Chromium single cell 3’ reagent kit v2 (10_X_ Genomics). cDNA libraries were sequenced on a Nextseq 500 (Illumina) to a minimum depth of 20,000 mean reads per cell. Demultiplexing, alignment to the modified mm10 reference genome, principal component analysis (PCA), clustering, non-linear reduction (t-SNE) and differential expression was performed using the cellranger count pipeline (10_X_ Genomics). Graphical output, merging and subsetting of datasets was performed using Seurat R package (Satija Lab, v.2.2). Subsetting was performed by excluding the clusters that expressed pericytic markers in order to focus only on the cFAPs when building a phylogenetic tree or merging the datasets. Subsequently, PCA, graph-based clustering, t-SNE and Wilcoxon rank sum test for differential expression were performed with Seurat R package. Gene ontology analysis for the 50 most differentially upregulated genes in the different clusters was performed using the DAVID Bioinformatics Resources (Huang et al., 2009a, 2009b) (v.6.8).

### Statistics

Power calculations were not done to predetermine sample size. We did not use any specific method of randomization to determine how samples and/or animals were allocated to experimental groups and processed. Analyses were not performed in a blinded fashion. No samples were excluded from the reported analyses. All results are presented as mean ± standard error of the mean (SEM). Statistical analysis was performed using unpaired Student’s t-test for analysis between 2 groups followed by Tukey’s post hoc test. For comparison between 3 or more groups, one-or two-way ANOVA with Bonferroni’s post-hoc test, with repeated measures in case of analyzing time course experiments, was performed using Graphpad Prism^®^ version 7 (GraphPad Software, La Jolla California USA). Sample size and/or replicate number for each experiment is indicated in the figure legends. Results with p values of less than 0.05 were considered statistically significant.

### Data availability

The bulk and single cell RNA sequencing datasets generated during the current study are available in the NCBI Gene Expression Omnibus (link will be provided upon manuscript acceptance).

