## Supplementary figures and images for "Pathogenic potential of Hic1 expressing cardiac stromal progenitors"

### Suppl. Fig.1

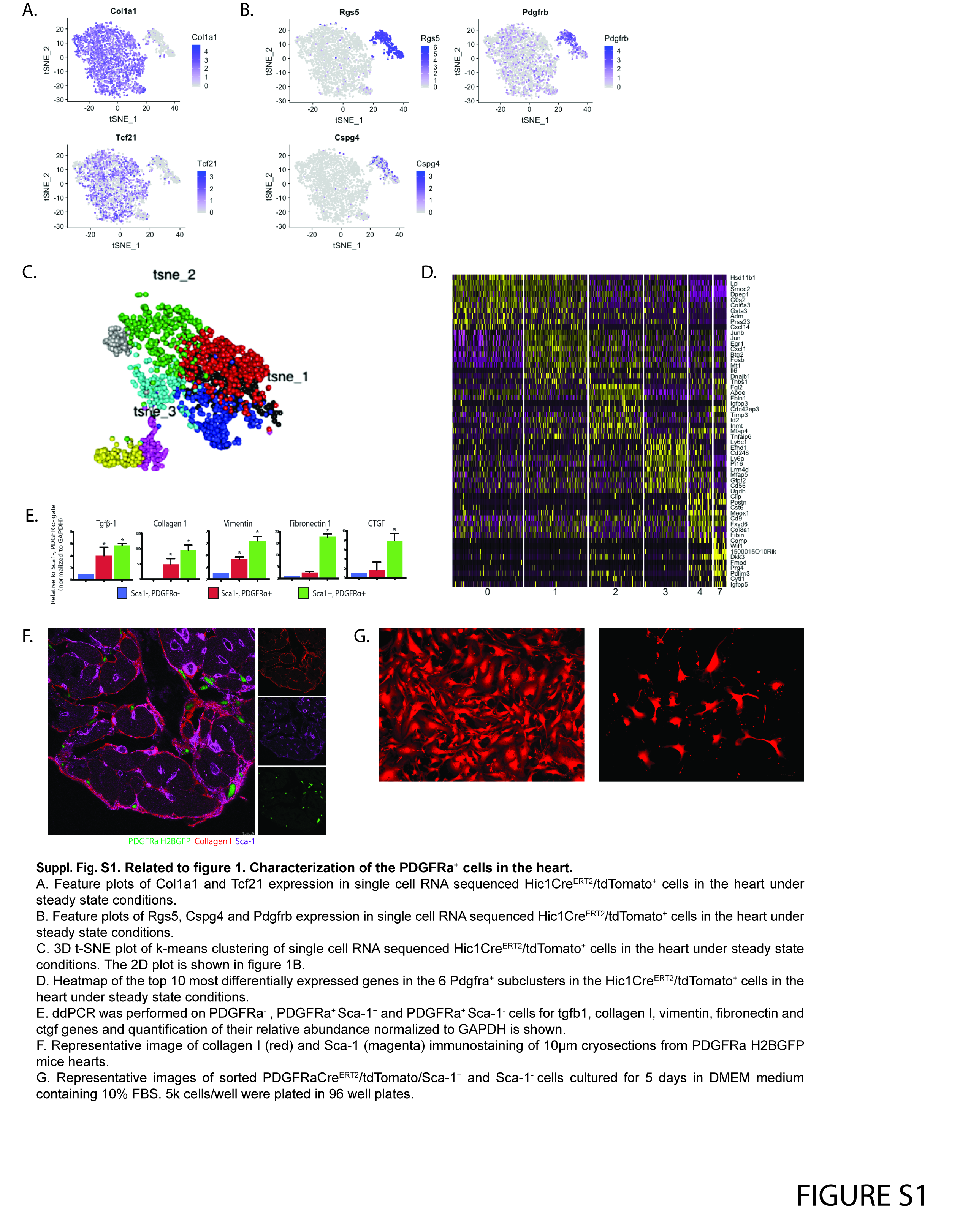

### Suppl. Fig.2

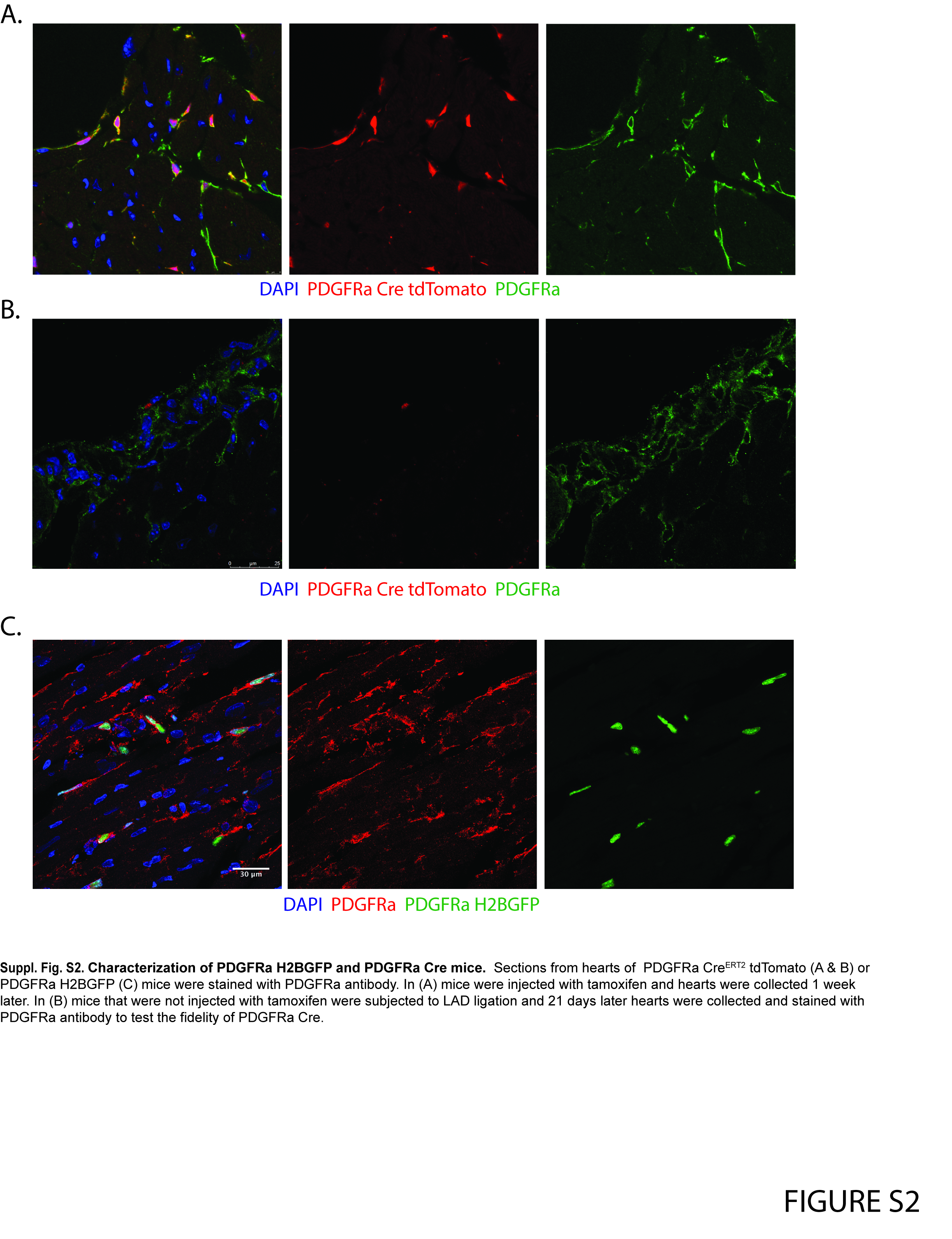

### Suppl. Fig.3

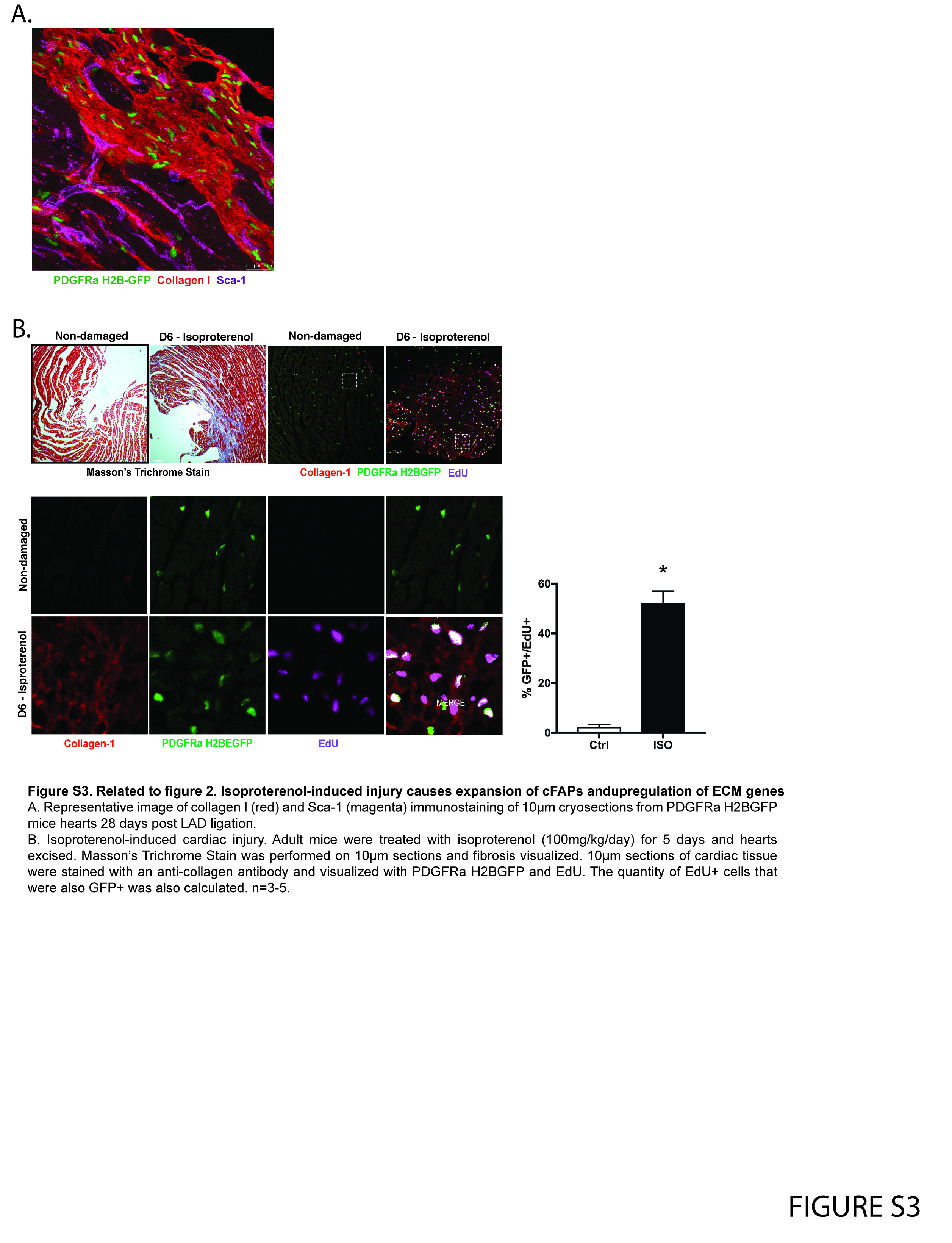

### Suppl. Fig.4

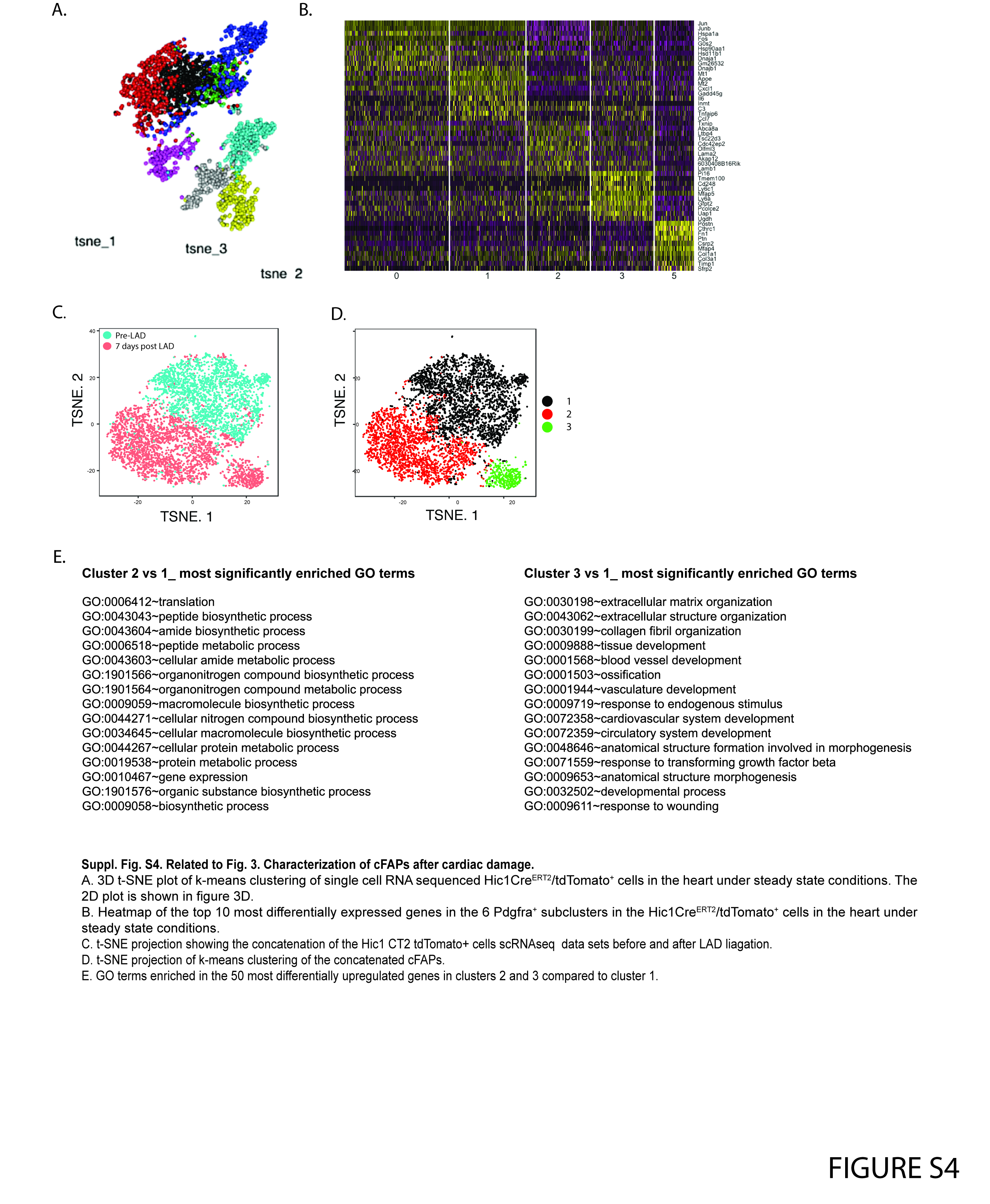

### Suppl. Fig. 5

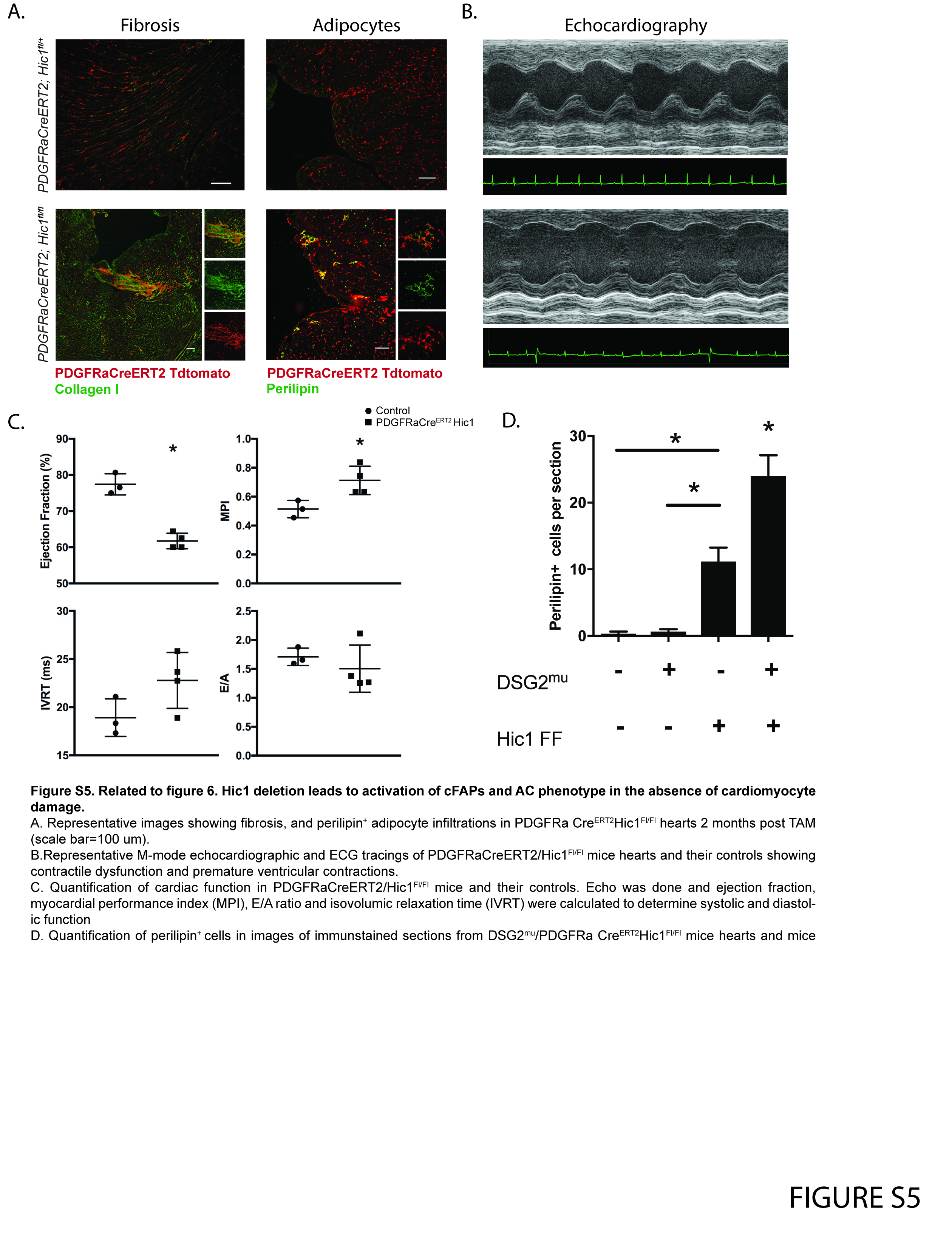
